# MicroEcoTools: an R package for comprehensive theoretical microbial ecology analysis

**DOI:** 10.1101/2024.08.19.608598

**Authors:** Soheil A. Neshat, Ezequiel Santillan, Stefan Wuertz

**Author notes:** Corresponding author: Ezequiel Santillan.

## Abstract

**Background:** Ecological theory aids in understanding how disturbances affect ecosystems. However, experimental data are often complex, with multiple post-disturbance theories potentially applying simultaneously to the same ecosystem. This emphasizes the need for tools to experimentally test these theoretical predictions.

**Results:** We introduce MicroEcoTools, an R package designed to test ecological framework predictions using microbial community data. It assesses microbial diversity and evaluates the relative impacts of stochastic and deterministic assembly mechanisms through a taxa-based null model approach for replicated designs. Specifically, the package allows application of Grime’s trait-based life-history categories—competitor, stress-tolerant, and ruderal (CSR)— to taxa, functional traits, and ecosystem functions within microbial communities. MicroEcoTools also includes relevant statistical tests, numeric simulations, and publicly available datasets for demonstration.

**Conclusions:** In conclusion, MicroEcoTools facilitates the application of ecological frameworks, including community assembly mechanisms, diversity analysis, and life-history strategies, to microbial ecosystems under disturbance. This R package, along with its source code, can be freely accessed on GitHub at https://www.github.com/Soheil-A-Neshat/MicroEcoTools.

## Background

Microbial ecology explores the intricate interactions and dynamics of microorganisms within ecosystems, spanning from soil^1^ and water^2^ to the human microbiome^3^. Understanding the complexity of microbial processes is crucial for elucidating ecosystem functioning, biogeochemical cycling^4^, and even human health^3^, especially under disturbance conditions^5^. Disturbances, ranging from natural events like fire and floods to anthropogenic ones such as releasing chemicals to water bodies, exert profound effects on ecosystems^6,7^. They can alter species richness, community composition, and functional traits, thereby impacting ecosystem processes^7–10^. The relationship between disturbance, diversity, and ecosystem function can be systematically explained by ecological theories^8,11^. These theories help to draw generalized conclusions from specific observations of organisms in their environment and thus classify, interpret, and predict the world around us^12^. To enable researchers to experimentally test predictions based on theory, new tools are needed to automate methods and streamline the analysis of large datasets.

Numerous theories examine how ecosystems respond to disturbances, and multiple post-disturbance theories may be applicable to the same ecosystem^13^. Several computational tools have been developed to analyse microbiome data with a strong focus on community structure, meaning the composition and arrangement of taxa within a community^14–17^. However, these tools often lack the capability to assess the relative contributions of stochastic and deterministic processes on community assembly. This is significant because community assembly processes are believed to shape community structure, thereby linking them to ecosystem function^9,18,19^. Recent developments in tools have begun to address these gaps^20,21^, yet additional advancements are needed to predict how microbial communities respond to environmental changes and disturbances across various theoretical frameworks.

Trait-based theory posits that organisms face tradeoffs when allocating resources to certain traits to maximize their fitness, which depend on abiotic and biotic interactions within their habitat^22,23^. The theoretical CSR framework for plant communities developed by Grime^24^ constitutes a classic trait-based approach and proposes three types of life-history strategies: competitors (C) who maximize resource acquisition and control in consistently productive niches, stress tolerants (S) who can maintain metabolic performance in unproductive niches, and ruderals (R) who have good growth rates but inefficient resource uptake in niches where events are frequently detrimental to the individual. Such strategies depend on varying intensities of disturbance (biomass destruction), stress (biomass restriction) and competition for resources. While this framework has been tested in microbial communities^23^, no computational tools are currently available to extend its application to broader studies.

We introduce MicroEcoTools, an R package designed to test ecological framework predictions using microbial community data. It assesses α-diversity and evaluates the relative impacts of stochastic and deterministic assembly mechanisms through a taxa-based null model analysis (NMA) across replicated designs. Additionally, MicroEcoTools extends traditional community ecology analyses to incorporate trait-based approaches, where traits denote functional characteristics influencing organisms’ ecological roles and responses to environmental changes. The package allows application of Grime’s trait-based framework categories—competitor, stress-tolerant, and ruderal (CSR)—to taxa, functional traits, and ecosystem functions within microbial communities. It includes relevant statistical tests, numeric simulations, and publicly available datasets for demonstration.

## Implementation

### Installation and usage

MicroEcoTools is an R package that depends on R ≥ 3.5 to run. The other dependencies in the R environment are data.table^25^, dplyr^26^, effectsize^27^, ggplot2^28^, magrittr^29^, plyr^26^, progress^30^, reshape2^31^, vegan^15^, parallel, doParallel^32^, and foreach^33^ packages. The MicroEcoTools package can be installed through the GitHub page (www.github.com/Soheil-A-Neshat/MicroEcoTools) using the install_github function of the devtools^30^ R package. MicroEcoTools contains two sets of functions for NMA and CSR related analysis (Table 1).

**TABLE 1.** The R functions imported by installing the MicroEcoTools package

| Category | Function | Description |
| --- | --- | --- |
| Null model analysis | NMA() | Takes in a dataset in long format and performs null model analysis comparing the observed communities and communities generated using the null assumption. |
| Trait-based analysis | Welch_ANOVA()<sup>†</sup> | Takes a dataset in wide format and performs a statistical test of means using the experimental group as explanatory variable, assuming non-constancy of variance. |
|  | pairwise_welch()<sup>†</sup> | Performs pairwise comparison on the data using the experimental group as explanatory variable, assuming non-constancy of variance. |
|  | CSR_assign() | Assigns competitor, stress-tolerant and ruderal categories to traits using the results of the pairwise comparison. |
|  | CSR_Simulation() | To predict the accuracy of the CSR assignments, this function utilizes a simulation algorithm based on the multinomial distribution to generate simulated communities and assigns competitor, stress-tolerant and ruderal categories to the observed and simulated communities. |
| Diversity analysis | Hill_diversity() | To calculate zero-, first- and second-order Hill numbers for the input count/relative abundance table. |

<sup>†</sup>Welch\_ANOVA and pairwise\_welch are utilised within CSR\_assign and CSR\_simulation functions, but they can also be called upon for custom statistical testing.

Detailed descriptions of the functions along with examples are available through the help function of the MicroEcoTools in the R environment.

### Simulation algorithms

The implemented simulation algorithm uses a multinomial distribution as the probability function to generate simulated count data. It takes in the observed community or traits data, which requires at least two replicates for a group (*i.e.*, minimum n = 2). Then, the algorithm generates simulated communities with the same total count of taxa or traits with the assumption that the probability for each individual to be present in any of the replicates is equal (Figure 1). For the CSR function, the algorithm calculates a probability distribution value from the observed data and generates simulated communities that have similar probability distributions.

**Figure 1.**
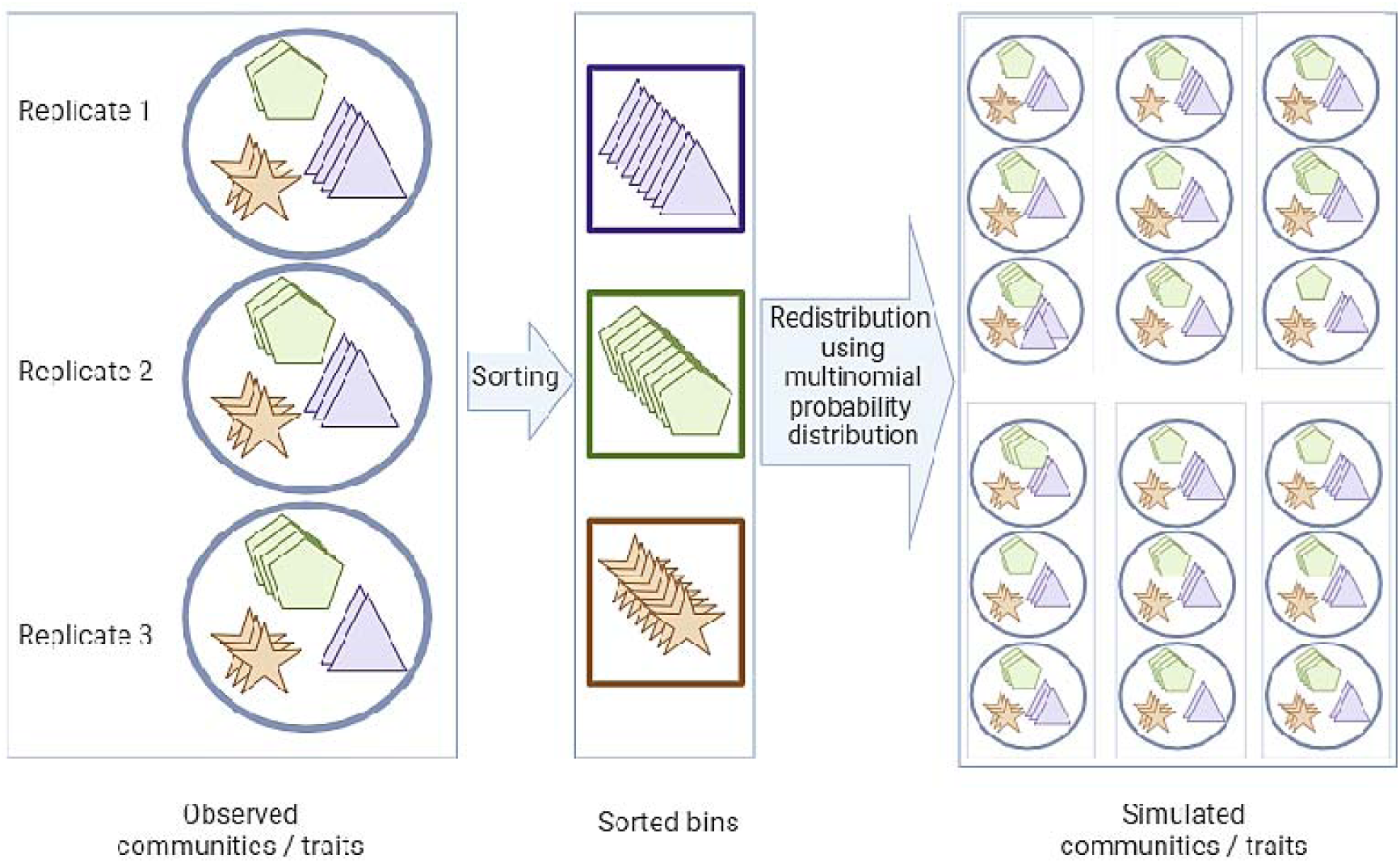
Schematic illustration of the simulation algorithm utilized by MicroEcoTools. This algorithm accepts count data as input and generates simulated count data using a multinomial distribution. Symbols within circles and squares represent different taxa or traits in the data.

### Null model analysis

As described by Nicholas Gotelli^34^, “null models are pattern generating models that deliberately exclude a mechanism of interest, and allow for randomization tests of ecological and biogeographic data”. The implemented null model analysis in the MicroEcoTools is adopted from a null model analysis tested on woody plants^35^ and wastewater treatment microbial communities^8^, with emphasis in flexibility and ease of use. This model uses the beta partition of alpha diversity using the following formula:

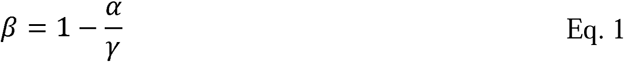

In this model, simulated communities are generated using the observed communities by randomly shuffling the species between samples (sites) while keeping the total number of observed species constant (Figure 1). After generating the simulated communities, diversity is measured using the vegan^15^ package for the observed and simulated communities (Figure 2). Then, the standard effect size (SES) is calculated as:

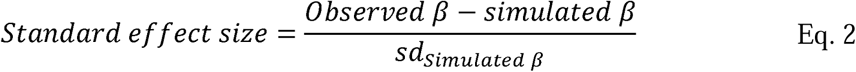

**Figure 2.**
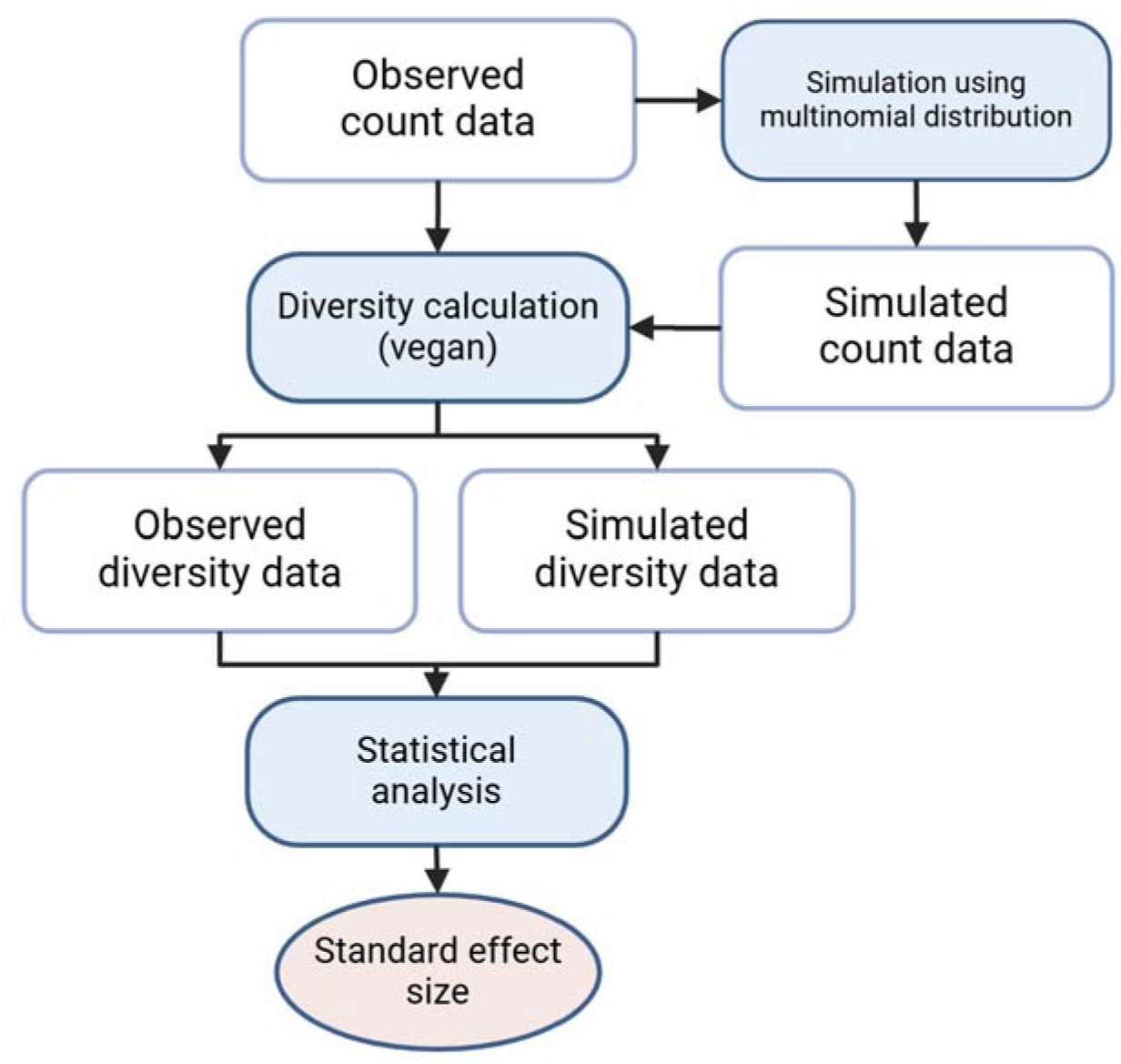
The null model analysis workflow implemented in the MicroEcoTools R package. The analysis is performed on count data with at least two biological replicates (minimum n = 2). The simulation is performed using the MicroEcoTools simulation algorithm while the diversity analysis is performed utilizing the vegan package.

The calculated SES represents the difference between communities formed under the influence of a mechanism of interest and simulated communities excluding the effect of that mechanism. The higher the SES, the greater the difference between those two communities, hence the higher the effect of deterministic community assembly. On the other hand, the lower the SES, the closer an observed community is to the null expectation, which is interpreted as a higher contribution of stochastic assembly mechanisms.

### Grime’s competitor – stress tolerant – ruderal (CSR) life-history category assignment

The Grime’s CSR framework^23,24,36^ classifies traits or community members into three main categories, namely, competitor, stress-tolerant and ruderals. Traits classified as competitor (C) are related to maximizing resource acquisition and control, while stress tolerant (S) and ruderal (R) traits are related to maintaining metabolic performance in high stress and high growth rate conditions, respectively^23^. The factors determining the class of a trait under the CSR framework are disturbance, stress, and competition for resources. Based on these factors, MicroEcoTools follows a decision tree approach (Figure 3), where traits abundant under low disturbance and high competition (undisturbed) conditions are classified as competitors (C), high disturbance and low stress (intermediately disturbed) conditions select for ruderals (R), and high stress and low disturbance (press-disturbed) conditions select for stress-tolerants (S). Composite classifications that arise from combining any of these three categories are also considered, being either CR, CS, SR, or CSR (Figure 3).

**Figure 3.**
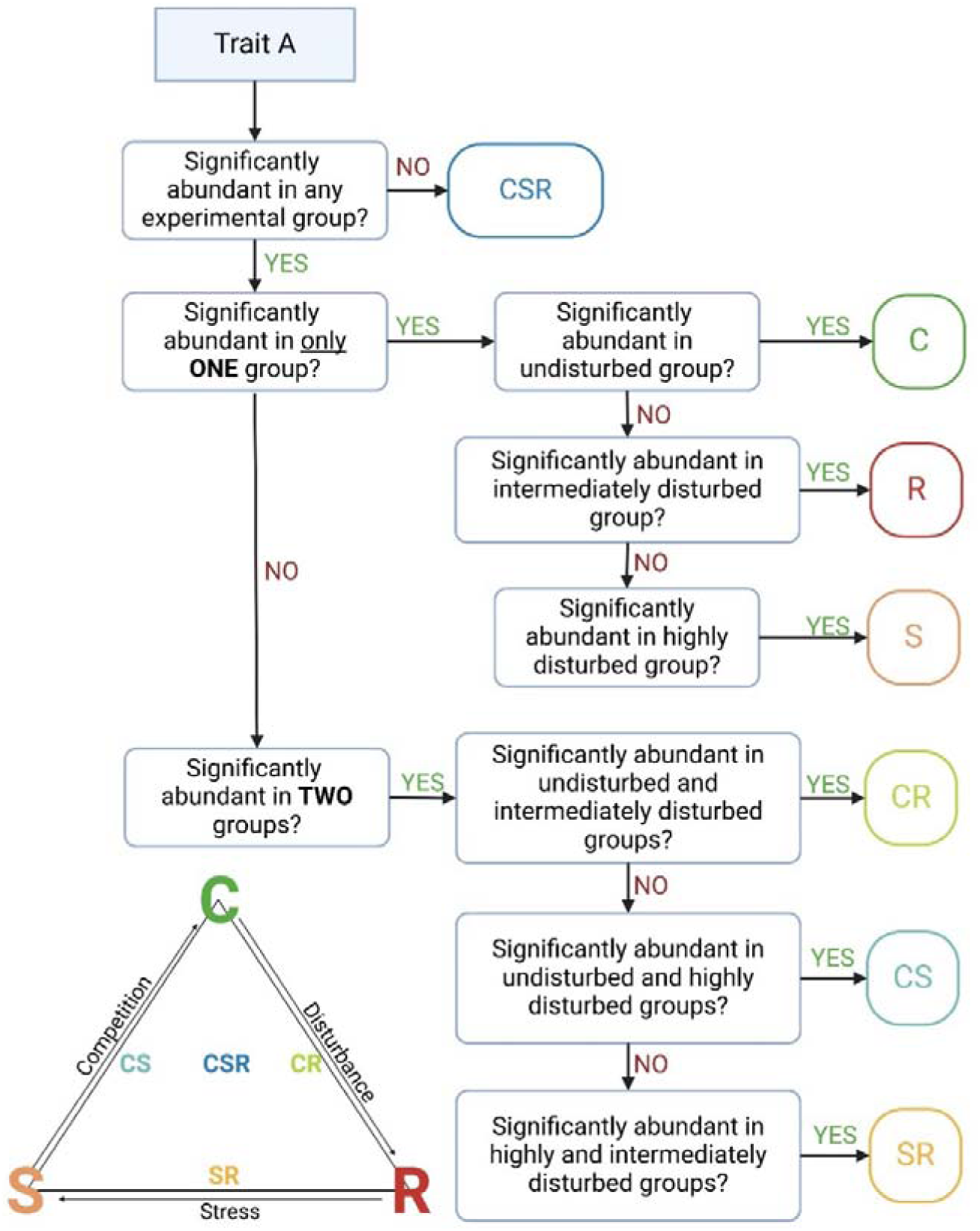
The Grime’s competitor (C), stress-tolerant (S), ruderal (R) framework and the decision tree utilized by the MicroEcoTools for classifying traits into these categories. The decision tree shows a scenario where there are only three groups in the experimental design, namely, undisturbed, intermediately disturbed and highly disturbed. The MicroEcoTools can also classify traits into the CSR categories in cases where there is more than one subgroup of the intermediately disturbed group. In that case, a trait will be categorized as C or S if it is significantly more abundant in undisturbed or highly disturbed groups compared to all intermediately disturbed subgroups, respectively. In case of an R trait, the trait should be significantly more abundant in any of the intermediately disturbed subgroups compared with undisturbed and highly disturbed groups. For more details on how traits are classified in more complex designs please visit the MicroEcoTools GitHub page at www.github.com/Soheil-A-Neshat/MicroEcoTools.

### Welch-ANOVA analysis

To test if a trait is significantly more abundant in one of the experimental settings, the MicroEcoTools uses the Welch-ANOVA test by default, given that non-constancy of variance is common for microbial datasets. This analysis assesses the statistical significance of the explanatory variable (disturbance level) explaining the changes in the response variable (community composition or abundance of functional traits). This function (MicroEcoTools::welch_ANOVA) is developed utilizing the one-way test function of the base R^30^. By default, this function assumes non-equality of variance between groups. The MicroEcoTools implementation of this function automatically detects if the variance in at least one group is zero. Upon detection of such an instance, it switches to ANOVA returning a warning in the remarks section of the generated results table.

### Pairwise Welch-ANOVA

The pairwise Welch-ANOVA function (MicroEcoTools::pairwise_welch) utilizes the combn function of the base R to generate all unique combinations of an input set. After generating the combination table consisting of all pairs of the elements in the experimental groups, the function uses the oneway.test function of the base R^30^ to assess the difference between two groups assuming non-equal variance. Similar to the MicroEcoTools Welch-ANOVA function, if the variance is zero in at least one group, it will switch to an ANOVA test generating a warning under the Remarks column in the generated report.

### MicroEcoTools datasets

The MicroEcoTools introduces two curated datasets that can be used to apply different ecological frameworks. These datasets are attached to the MicroEcoTools as dataframes under the names NMA_data^8^ and CSR_data^23^. The datasets were generated in prior studies described by Santillan et al.^8,23^ and can be imported into R using the data function of the base R^30^. The structure of input data should follow the structure of these datasets for the MicroEcoTools functions to run smoothly.

### Experimental design for the datasets included in the MicroEcoTools

The datasets were generated in a previous experiment employing sequencing batch microcosm bioreactors (20-mL working volume) inoculated with activated sludge from a full-scale plant and operated for 35 days. These reactors were fed daily using a complex synthetic feed. 3-chloroaniline was added to the feed as disturbance at varied frequencies during the experiment. The experiment involved eight levels of disturbance in triplicate independent reactors (n = 24). These reactors were subjected to 3-CA either daily (press-disturbed), every 2, 3, 4, 5, 6, or 7 days (intermediately disturbed), or left undisturbed. Each level was assigned a numerical value ranging from 0 to 7. Details of the sample preparation, sequencing, and bioinformatics pipelines are explained in Santillan et al.^8,23^.

## Results

To demonstrate the functions implemented in the MicroEcoTools package, we included two datasets, NMA_data and CSR_data, which are publicly available^37,38^. Briefly, the NMA_data dataset contains microbial community genus-level count data, and the CSR_data dataset contains functional category abundance data obtained in a study where bacterial communities where exposed to different frequencies of disturbance^8,23^. These datasets can be imported into R to explore the data structure and test the implemented functions.

### Example 1. NMA_data dataset: effect of disturbance frequency on assembly mechanisms in activated sludge bioreactors

The NMA_data dataset was generated in a disturbance experiment that led to the intermediate stochasticity hypothesis (ISH)^8^, based on a test of the intermediate disturbance hypothesis (IDH)^11^. The ISH poses that the peak in α-diversity observed at intermediate levels of disturbance has a mechanistic explanation due to a shift in the relative contribution of stochastic and deterministic assembly mechanisms. To test this hypothesis, after observing the peak in α-diversity at intermediate disturbance levels predicted by the IDH (Figure 4a), a taxa-based null model analysis was performed using the NMA function on the count NMA_data using taxa richness as the α-diversity metric to calculate the β-partition, as in previous studies^8,39^. In the case of relative abundance data provided by the user, the function converts it to counts by multiplying it by 100,000 before applying the necessary simulations. The advantages of using MicroEcoTools for this task are twofold. First, the simulation algorithm is computationally more efficient than before^8^. Second, the NMA function also allows the user to set zero, first, or second order Hill numbers^40^ as measures of α-diversity for the null model analysis. The output of the analysis is a standard effect size (SES) plot (Figure 4b). In the case of this dataset, the SES is lower at intermediate disturbance frequency levels, meaning that communities in such reactors are closer to the null expectation than at other levels at the low or high end of the frequency range tested. This is interpreted as a higher contribution of stochastic assembly mechanisms at intermediate levels of disturbance, in accordance with the ISH ^8,9^. Note that different patterns in SES could arise when using different orders of Hill α-diversity; the assessment depends on the type of data available, and the question being addressed in each study. Also, the values of SES are specific to each dataset and are not directly comparable across studies. Finally, as all replicates per group are used to calculate the β-partition as in Eq. 1, one SES value per group is calculated.

**Figure 4.**
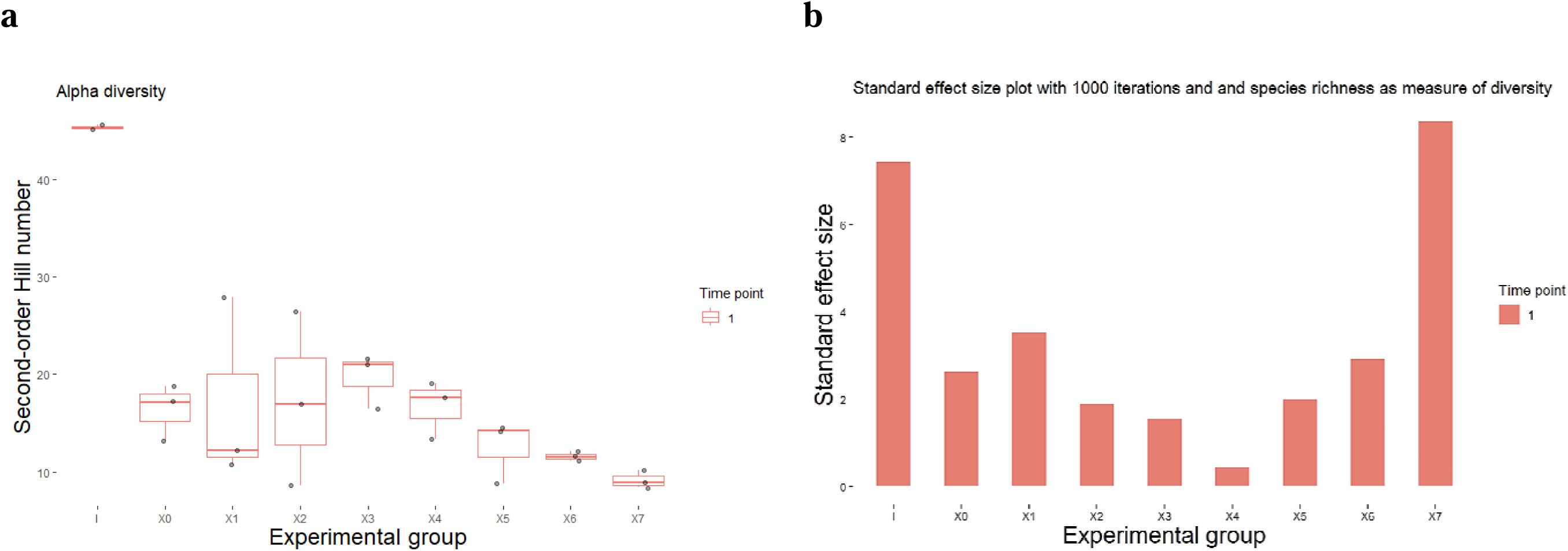
Output plots generated by the MicroEcoTools package. (**a**) Diversity and (**b**) standard effect size plots were generated by the NMA function with 1000 simulations applied to the example datasets included in the MicroEcoTools package. Community data was generated using metabarcoding. The x-axis shows the experimental groups (I: inoculum, X0-X7: disturbance range from no disturbance to press disturbance) and the y-axis the standard effect size generated by comparing communities generated will a null hypothesis and observed communities.

### Example 2. CSR_data dataset: competitor, stress-tolerant, and ruderal life-history strategies adopted by undisturbed, press disturbed and intermediately disturbed communities, respectively

A balanced replicated experimental design allowed us to apply the CSR framework to the aggregated traits data generated by Santillan et al.^23^. In that study^23^, the reactors were exposed to three main regimes of disturbance: undisturbed, intermediately disturbed, and press disturbed. Through multivariate statistical analysis, the authors demonstrated a three-way separation in both taxonomic and genotypic trait metagenomics data, suggesting that the communities under different regimes of disturbance frequency adopted different life history strategies. However, the process of examining each taxon and functional trait in their study was manual and laborious. The CSR_assign function of MicroEcoTools allows for the automatic classification of taxa and traits under the CSR framework based on a decision tree (Figure 3), enabling the plotting of the most abundant traits from each life-history category, including all possible combinations: C, S, R, CS, CR, and SR (Figure 5).

**Figure 5.**
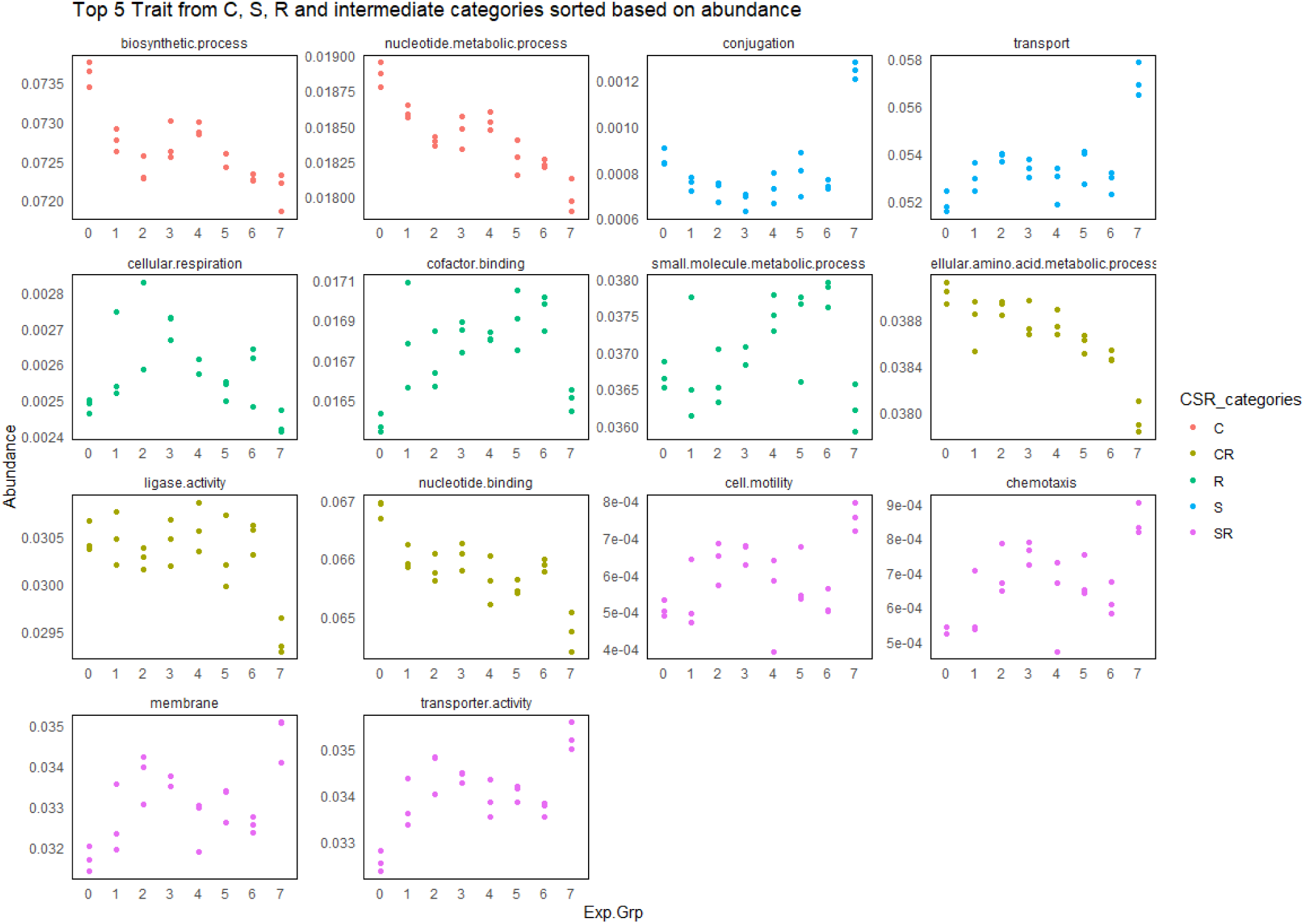
Grime’s CSR category assignment using the CSR assign function of the MicroEcoTools on the traits annotated with the IP2G^41^ database. The x-axis shows the experimental groups (Exp.Grp) 0 to 7 representing a range of disturbances from no disturbance (0) to press disturbance (7), with intermediate levels of disturbance in groups 1 to 6. The y-axis represents the abundance of functional traits. The color of data points on each panel refers to different assigned CSR categories.

We observed that, in some cases, a slight change in the count data can alter the type of CSR assignments, highlighting the importance of replication in experimental designs. Because replication is not always feasible due to various constrains, such as limited resources, we implemented a simulation-based approach in MicroEcoTools to estimate the accuracy of CSR category assignments. This approach leverages the MicroEcoTools simulation algorithm, which assumes that the same probability distribution will be observed if the experiment is repeated (Figure 1). The algorithm generates simulated communities and runs the CSR assignment function for all the simulated communities. The resultant dataset is used to estimate the likelihood of a trait being assigned to a CSR category. Table 2 shows the results of the analysis on the simulated genotypic trait data, predicting the accuracy of the CSR assignments.

**Table 2.** The first 15 rows in alphabetic order from the CSR_assign function output, generated with 1000 simulations to assess the accuracy of the CSR category assignments. The first column shows the trait the analysis was performed on, while columns 2 to 6 display the output of the Welch-ANOVA function of MicroEcoTools, revealing the statistical comparison between experimental groups.

| Trait | Welch-ANOVA p-value<sup>1</sup> | Method | Adjusted Welch-ANOVA p-value<sup>2</sup> | CSR categories | Remarks | No. of times a trait is assigned to a CSR category in simulated communities (%)<sup>2</sup> |  |  |  |  |  |  |  |
| --- | --- | --- | --- | --- | --- | --- | --- | --- | --- | --- | --- | --- | --- |
|  |  |  |  |  |  | C | R | S | CR | CS | SR | CSR | NA |
| ATP.binding.cassette..ABC..transporter.complex | 0.00289 | One-way analysis of means (not assuming equal variances) | 0.00521 | CSR | - | 0.00 | 0.00 | 0.90 | 0.00 | 0.40 | 0.10 | 98.60 | 0.00 |
| biosynthetic.process | 0.00020 | One-way analysis of means (not assuming equal variances) | 0.00158 | C | - | 0.20 | 0.00 | 0.00 | 0.50 | 0.00 | 0.00 | 99.30 | 0.00 |
| carbohydrate.metabolic.process | 0.00123 | One-way analysis of means (not assuming equal variances) | 0.00334 | CSR | - | 0.00 | 0.30 | 0.00 | 5.89 | 0.00 | 0.00 | 93.81 | 0.00 |
| catabolic.process | 0.00002 | One-way analysis of means (not assuming equal variances) | 0.00104 | CSR | - | 0.00 | 0.80 | 0.00 | 0.00 | 0.00 | 0.50 | 98.70 | 0.00 |
| cell.communication | 0.10141 | One-way analysis of means (not assuming equal variances) | 0.10510 | CSR | - | 0.00 | 0.00 | 0.00 | 0.10 | 0.00 | 0.00 | 99.90 | 0.00 |
| cell.division | 0.00705 | One-way analysis of means (not assuming equal variances) | 0.01057 | CSR | - | 0.00 | 0.00 | 0.00 | 0.00 | 0.10 | 0.00 | 99.90 | 0.00 |
| cell.motility | 0.00293 | One-way analysis of means (not assuming equal variances) | 0.00521 | SR | - | 0.00 | 0.00 | 0.00 | 0.00 | 0.00 | 1.80 | 98.20 | 0.00 |
| cell.wall | 0.00249 | One-way analysis of means (not assuming equal variances) | 0.00489 | CSR | - | 0.00 | 0.00 | 0.00 | 0.60 | 0.00 | 0.00 | 99.40 | 0.00 |
| cellular.amino.acid.metabolic.process | 0.00051 | One-way analysis of means (not assuming equal variances) | 0.00211 | CR | - | 0.00 | 0.00 | 0.00 | 1.10 | 0.00 | 0.00 | 98.90 | 0.00 |
| cellular.component.organization | 0.00057 | One-way analysis of means (not assuming equal variances) | 0.00215 | CSR | - | 0.00 | 0.00 | 0.00 | 0.00 | 0.00 | 0.00 | 100.00 | 0.00 |
| cellular.respiration | 0.00192 | One-way analysis of means (not assuming equal variances) | 0.00406 | R | - | 0.00 | 1.00 | 0.00 | 0.10 | 0.00 | 0.00 | 98.90 | 0.00 |
| chemotaxis | 0.00043 | One-way analysis of means (not assuming equal variances) | 0.00206 | SR | - | 0.00 | 0.00 | 0.00 | 0.00 | 0.00 | 4.20 | 95.80 | 0.00 |
| chromosome | 0.05021 | One-way analysis of means (not assuming equal variances) | 0.05504 | CSR | - | 0.00 | 0.00 | 0.00 | 0.30 | 0.00 | 0.00 | 99.70 | 0.00 |
| cofactor.binding | 0.00029 | One-way analysis of means (not assuming equal variances) | 0.00158 | R | - | 0.00 | 1.00 | 0.00 | 0.00 | 0.00 | 0.10 | 98.90 | 0.00 |

<sup>1</sup> P-values are rounded to show 5 decimal places
<sup>2</sup> Assuming the same probability distribution as for the observed community. The last column shows the percentage of times the algorithm was unable to assign a category to a trait.

## Discussion

The MicroEcoTools package is designed to address computational gaps for microbial ecologists applying ecological frameworks. The algorithms enable users to test ecological theory on experimentally acquired data and are designed to provide a fast and seamless experience for those with basic R proficiency by minimizing the amount of coding required. Additionally, we implemented extensive error reporting functionalities to further enhance the user experience. Researchers often encounter issues related to data structure and experimental design when using R packages. To address this, MicroEcoTools runs pre-analysis assessments to check data quality, such as data structure and number of replicates, and returns detailed error messages. These messages include instructions to help users rectify the issues. Furthermore, MicroEcoTools can store data objects containing intermediate results generated at different workflow steps. This functionality has two main benefits: it helps users understand how complex results were generated and saves the simulated data for use in other analyses.

Another major gap in microbial ecology analyses that MicroEcoTools addresses is the limited availability of simulation algorithms. Most available tools that run null model analyses require generating communities with null assumptions, but the simulated communities are often purged after each iteration for efficiency considerations. MicroEcoTools uses a list object to store the simulated communities with different assumptions, which can be used subsequently for troubleshooting and as input for other analyses. Additionally, this function is very useful for teaching purposes, allowing the generation of different communities with varying probability distributions.

Lastly, accuracy in coding, correctness of results, reproducibility, and computational efficiency were prioritized in the development of MicroEcoTools. We tested the algorithms in MicroEcoTools on datasets from different ecosystems, including activated sludge, anaerobic digestion, and drinking water, and meticulously investigated the results to ensure accuracy. This process helps to ensure that the generated results are sound and reliable. Inaccurate results from computational tools can alter ecological interpretations, leading to misleading conclusions. Additionally, we validated the results generated by MicroEcoTools through cross-comparisons with the publications from which the NMA_data and CSR_data datasets were obtained^8,23^. This analysis showed that MicroEcoTools can produce accurate results that agree with and improve upon previous analyses, all while reducing analysis time and minimizing human error from manual curation.

## Conclusions

Here, we introduce the MicroEcoTools, an R package for applying ecological frameworks on microbial ecosystems. The current version of MicroEcoTools can apply a taxa-based null model analysis to assess relative contribution of stochastic and deterministic mechanisms in the community assembly and assign Grime’s competitor, stress-tolerant and ruderals (CSR) life-history strategy categories to taxa or genotypic traits. This life-history strategies framework simplifies complex datasets encompassing microbial traits, function, and taxa into ecologically meaningful components. MicroEcoTools benefits from a computationally efficient simulation algorithm implemented to analysing multidimensional community data. This package can help enhance our understanding of the response of microbial ecosystems to disturbance, allow testing of different theoretical frameworks, and aid into the development of new theories.

## Availability and requirements

Project name: MicroEcoTools

Project web page: https://www.github.com/Soheil-A-Neshat/MicroEcoTools

Official/formal distribution: https://www.github.com/Soheil-A-Neshat/MicroEcoTools

Help documentation: https://www.github.com/Soheil-A-Neshat/MicroEcoTools/tree/main/man

Operating system(s): Tested on R running on Linux CentOS 7, Apple Mac OS X Sonoma v14.3.1, and Microsoft Windows 11 v22H2 (OS Build 22261.3155)

Programming language: R

License: GNU GPLv3

## Declarations

The authors declare no conflicts of interest.

## Code availability

The codes are available through the MicroEcoTools Github page https://www.GitHub.com/Soheil-A-Neshat/MicroEcoTools.

## Limitations

The analysis can only be performed on data with at least two independent biological replicates. MicroEcoTools assesses the data before starting the main workflow to check the number of replicates and returns an error if any of the experimental groups are not meeting this requirement. In addition, the number of simulations can affect the results generated by the algorithms. Although the user can choose for any number of simulations, we cannot guarantee the accuracy of results generated with less than 1,000 simulations.

## Authors’ contributions

SN, ES and SW conceived the study. SN and ES developed the algorithms. SN carried out the coding and R implementation. SN and ES performed test runs. SW secured the funding for the project. SN and ES wrote the first manuscript draft. All authors contributed to writing and reviewing the manuscript.

## Acknowledgements

This project was supported by the Singapore National Research Foundation and Ministry of Education of Singapore under the Research Centre of Excellence Program.

## References

1. Paul, E. & Frey, S. Soil Microbiology, Ecology and Biochemistry. (Elsevier, 2023).

2. Grossart, H., Massana, R., McMahon, K. D. & Walsh, D. A. Linking metagenomics to aquatic microbial ecology and biogeochemical cycles. Limnology and Oceanography 65, S2–S20 (2020).

3. Ogunrinola, G. A., Oyewale, J. O., Oshamika, O. O. & Olasehinde, G. I. The human microbiome and its impacts on health. International journal of microbiology 2020, (2020).

4. Kappler, A. et al. An evolving view on biogeochemical cycling of iron. Nature Reviews Microbiology 19, 360–374 (2021).

5. Neshat, S., Santillan, E., Seshan, H. & Wuertz, S. Non-redundant metagenome-assembled genomes of activated sludge reactors at different disturbances and scales. Scientific Data 11, 855 (2024).

6. Korpinen, S. et al. Combined effects of human pressures on Europe’s marine ecosystems. Ambio 50, 1325–1336 (2021).

7. Battisti, C., Poeta, G. & Fanelli, G. The Concept of Disturbance BT - An Introduction to Disturbance Ecology: A Road Map for Wildlife Management and Conservation. in (eds. Battisti, C., Poeta, G. & Fanelli, G.) 7–12 (Springer International Publishing, Cham, 2016). doi:10.1007/978-3-319-32476-0_2.

8. Santillan, E., Seshan, H., Constancias, F., Drautz-Moses, D. I. & Wuertz, S. Frequency of disturbance alters diversity, function, and underlying assembly mechanisms of complex bacterial communities. npj Biofilms and Microbiomes 5, 1–8 (2019).

9. Santillan, E. & Wuertz, S. Microbiome assembly predictably shapes diversity across a range of disturbance frequencies in experimental microcosms. npj Biofilms and Microbiomes 8, 1–11 (2022).

10. Neshat, S. Microbiome Studies on Anaerobic Digestion Using Genome–Resolved Multi– Omics. (Nanyang Technological University Singapore, Singapore, 2022). doi:10.32657/10356/168304.

11. Connell, J. H. Diversity in tropical rain forests and coral reefs. Science 199, 1302–1310 (1978).

12. Prosser, J. I. et al. The role of ecological theory in microbial ecology. Nature Reviews Microbiology 5, 384–384 (2007).

13. Pulsford, S. A., Lindenmayer, D. B. & Driscoll, D. A. A succession of theories: purging redundancy from disturbance theory. Biological Reviews 91, 148–167 (2016).

14. Schloss, P. D. Reintroducing mothur: 10 years later. Applied and environmental microbiology 86, e02343–19 (2020).

15. Jari Oksanen, F. G. B., Michael Friendly, Roeland Kindt, Pierre Legendre, Dan McGlinn, P. R. M., R. B. O’Hara, Gavin L. Simpson, Peter Solymos, M. Henry & H. Stevens, E. S. and H. W. vegan: Community Ecology Package. R. (2019).

16. Bolyen, E. et al. Reproducible, interactive, scalable and extensible microbiome data science using QIIME 2. Nature Biotechnology 37, 852–857 (2019).

17. McMurdie, P. J. & Holmes, S. phyloseq: an R package for reproducible interactive analysis and graphics of microbiome census data. PloS one 8, e61217 (2013).

18. Santillan, E., Constancias, F. & Wuertz, S. Press Disturbance Alters Community Structure and Assembly Mechanisms of Bacterial Taxa and Functional Genes in Mesocosm-Scale Bioreactors. mSystems 5, 1–20 (2020).

19. Santillan, E., Seshan, H. & Wuertz, S. Press Xenobiotic 3-Chloroaniline Disturbance Favors Deterministic Assembly with a Shift in Function and Structure of Bacterial Communities in Sludge Bioreactors. ACS ES&T Water 1, 1429–1437 (2021).

20. Ning, D. et al. A quantitative framework reveals ecological drivers of grassland microbial community assembly in response to warming. Nature Communications 11, 4717 (2020).

21. Liu, C., Cui, Y., Li, X. & Yao, M. microeco: an R package for data mining in microbial community ecology. FEMS microbiology ecology 97, fiaa255 (2021).

22. Krause, S. Trait-based approaches for understanding microbial biodiversity and ecosystem functioning. Frontiers in Microbiology 5, 1–10 (2014).

23. Santillan, E., Seshan, H., Constancias, F. & Wuertz, S. Trait-based life-history strategies explain succession scenario for complex bacterial communities under varying disturbance. Environmental Microbiology 21, 3751–3764 (2019).

24. Grime, J. P. Evidence for the existence of three primary strategies in plants and its relevance to ecological and evolutionary theory. The american naturalist 111, 1169–1194 (1977).

25. Dowle, M., et al. Package ‘data. table’. Extension of ‘data. frame 596, (2019).

26. Wickham, H., Francois, R., Henry, L. & Müller, K. dplyr: A grammar of data manipulation. R package version 0.4 (2015).

27. Ben-Shachar, M. S., Lüdecke, D. & Makowski, D. effectsize: Estimation of effect size indices and standardized parameters. Journal of Open Source Software 5, 2815 (2020).

28. Wickham, H. ggplot2. Wiley Interdisciplinary Reviews: Computational Statistics 3, 180– 185 (2011).

29. Wickham, H. et al. Welcome to the Tidyverse. Journal of Open Source Software 4, 1686– 1686 (2019).

30. R Core Team. R: A language and environment for statistical computing. (2018).

31. Wickham, H. reshape2: Flexibly reshape data: a reboot of the reshape package. R Package version 1.2 (2012).

32. Calaway, R., Analytics, R., Weston, S., Tenenbaum, D. & Calaway, M. R. Package “doParallel”. (2015).

33. Weston, S. Using the foreach package. (2019).

34. Gotelli, N. J. Research frontiers in null model analysis. Global Ecology and Biogeography 10, 337–343 (2001).

35. Kraft, N. J. B. Disentangling the drivers of β diversity along latitudinal and elevational gradients. Science 333, 1755–1758 (2011).

36. Grime, J. P. & Pierce, S. The Evolutionary Strategies That Shape Ecosystems. (John Wiley & Sons, 2012).

37. Santillan, E., Seshan, H., Constancias, F., Drautz-Moses, D. & Wuertz, S. Sludge bioreactor microcosm study on complex bacterial communities and ecosystem function using 3-CA disturbance at varying frequencies for 35 days. (Figshare dataset). Figshare 10.6084/m9.figshare.7369964.v1 (2019).

38. Santillan, E., Seshan, Ha., Constancias, F. & Wuertz, S. Trait-based life-history strategies study on complex bacterial communities ecosystem function and succession using 3-CA disturbance at varying frequencies for 35 days in sludge bioreactors (Figshare dataset). Figshare 10.6084/m9.figshare.7967480.v1 (2019).

39. Kraft, N. J. B. Disentangling the drivers of β diversity along latitudinal and elevational gradients. Science 333, 1755–1758 (2011).

40. Hill, M. O. Diversity and evenness: a unifiying notation and its consequences. Ecology 54, 427–432 (1973).

41. Paysan-Lafosse, T. et al. InterPro in 2022. Nucleic acids research 51, D418–D427 (2023).

